# *Streptococcus agalactiae* induces placental macrophages to release extracellular traps loaded with tissue remodeling enzymes via an oxidative-burst-dependent mechanism

**DOI:** 10.1101/440685

**Authors:** Ryan S. Doster, Jessica A. Sutton, Lisa M. Rogers, David M. Aronoff, Jennifer A. Gaddy

## Abstract

*Streptococcus agalactiae*, or Group B *Streptococcus* (GBS), is a common perinatal pathogen. GBS colonization of the vaginal mucosa during pregnancy is a risk factor for invasive infection of the fetal membranes (chorioamnionitis) and its consequences such as membrane rupture, preterm labor, stillbirth, and neonatal sepsis. Placental macrophages, or Hofbauer cells, are fetally-derived macrophages present within placental and fetal membrane tissues that perform vital functions for fetal and placental development, including supporting angiogenesis, tissue remodeling, and regulation of maternal-fetal tolerance. Although placental macrophages, as tissue-resident innate phagocytes, are likely to engage invasive bacteria such as GBS, there is limited information regarding how these cells respond to bacterial infection. Here, we demonstrate *in vitro* that placental macrophages release macrophage extracellular traps (METs) in response to bacterial infection. Placental macrophage METs contain proteins including histones, myeloperoxidase, and neutrophil elastase similar to neutrophil extracellular traps and are capable of killing GBS cells. MET release from these cells occurs by a process that depends on the production of reactive oxygen species. Placental macrophage METs also contain matrix metalloproteases that are released in response to GBS and could contribute to fetal membrane weakening during infection. MET structures were identified within human fetal membrane tissues infected *ex vivo*, suggesting that placental macrophages release METs in response to bacterial infection during chorioamnionitis.

**Importance:** *Streptococcus agalactiae*, also known as Group B *Streptococcus* (GBS), is a common pathogen during pregnancy where infection can result in chorioamnionitis, preterm premature rupture of membranes (PPROM), preterm labor, stillbirth, and neonatal sepsis. Mechanisms by which GBS infection results in adverse pregnancy outcomes are still incompletely understood. This study evaluated interactions between GBS and placental macrophages. The data demonstrate that in response to infection, placental macrophages release extracellular traps capable of killing GBS. Additionally, this work establishes that proteins associated with extracellular trap fibers include several matrix metalloproteinases that have been associated with chorioamnionitis. In the context of pregnancy, placental macrophage responses to bacterial infection might have beneficial and adverse consequences, including protective effects against bacterial invasion but also releasing important mediators of membrane breakdown that could contribute to membrane rupture or preterm labor.

## Introduction

15 million cases of preterm birth, or birth before 37 weeks gestation, occur annually worldwide, including 500,000 cases in the United States, conferring an estimated cost of $26.2 billion (1–3). The World Health Organization estimates that preterm birth complications are a leading cause of death among children under five years of age, resulting in nearly 1 million deaths in 2015 (4, 5). In addition to loss of child lives, preterm birth increases risk of chronic health conditions including neurodevelopmental deficits, metabolic syndrome, cardiovascular abnormalities, chronic kidney disease, and chronic respiratory conditions (6, 7).

*Streptococcus agalactiae*, also known as Group B *Streptococcus* (GBS), is a common perinatal pathogen (8). Approximately 10-40% of women are colonized with GBS during late pregnancy (9, 10). Rectovaginal GBS carriage is associated with adverse pregnancy outcomes including stillbirth, preterm labor, chorioamnionitis, and neonatal sepsis (11–13). Because of the burden and severity of GBS-related adverse pregnancy outcomes, the CDC recommends GBS screening late in gestation and antibiotic prophylaxis during labor (14). This strategy has decreased the incidence of early-onset neonatal sepsis but misses mothers that deliver preterm, before screening is conducted (14). Despite screening and treatment interventions, GBS remains a leading neonatal pathogen (15).

Pregnancy represents a unique immunologic state in which the maternal immune system must dampen its responses against foreign antigens of the semiallogenic fetus while defending the gravid uterus from infection. Excessive inflammation can drive adverse pregnancy events including loss of pregnancy, preterm birth, intrauterine growth restriction, and preeclampsia (16). Multiple mechanisms exist to support maternal-fetal tolerance including production of anti-inflammatory cytokines that alter the number and function of immune cells at the maternal-fetal interface (17–19). Unfortunately, infection is a common complication of pregnancy. Bacterial infection of the fetal membranes, known as chorioamnionitis, occurs most often by ascending infection from the vagina (8, 20, 21). During infection, bacterial products are recognized by pathogen recognition receptors, which then stimulate production of proinflammatory cytokines (20, 22, 23). These inflammatory mediators initiate a cascade of events that result in neutrophil infiltration into the fetal membranes, production and release of matrix metalloproteases (MMPs), and cervical contractions which eventually result in membrane rupture and preterm birth (24).

Macrophages represent 20-30% of the leukocytes within gestational tissues (25). In particular, fetally-derived macrophages, called Hofbauer cells or placental macrophages (PMs), play key roles in placental invasion, angiogenesis, tissue remodeling, and development (26, 27). The inflammatory state of these cells is carefully regulated throughout pregnancy. As the pregnancy progresses the M2 or anti-inflammatory and tissue remodeling phenotype predominates to supports fetal development (28–31). PMs contribute to immune tolerance by secretion of anti-inflammatory cytokines, which suppress production of proinflammatory cytokines (32–35). Disruption of appropriate macrophage polarization is associated with abnormal pregnancies including spontaneous abortions, preterm labor, and preeclampsia (28). We sought to understand how bacterial infection alters PM functions, and how these responses may contribute to pathologic pregnancies. These studies demonstrate that both PMs and a model macrophage cell line, the PMA-differentiated THP-1 macrophage-like cells, release macrophage extracellular traps (METs) in response to bacterial infection in a process that is dependent upon the generation of reactive oxygen species (ROS). METs, reminiscent of neutrophil extracellular traps (NETs), have recently been recognized as structures released by macrophages under a number of conditions including infection (36). PM METs contain histones, myeloperoxidase, and neutrophils elastase as well as several MMPs, and MET structures are found within human fetal membranes infected with GBS *ex vivo*.

## Results

### Placental macrophages release METs in response to GBS

To understand PM responses to GBS at the host-pathogen interface, isolated PMs were infected *ex vivo* with GBS and cellular interactions examined using field-gun high-resolution scanning electron microscopy (SEM). At one hour following infection, fine, reticular structures were noted extending from macrophages, and these structures were less abundant in uninfected samples (Figure 1A, lower panels). These structures resembled NETs. Recent reports suggest that macrophages also release fibers composed of DNA and histones, known as METs (36, 37). To determine if these structures were METs, macrophages were evaluated by scanning laser confocal microscopy after staining with the DNA binding dye SYTOX Green, which demonstrated extracellular structures extending from PMs that were not seen when PMs were treated with DNase I (Figure 1A, top panels). Cells were then evaluated to assess the degree to which these structures contained proteins previously associated with NETs and METs, including histones, myeloperoxidase, and neutrophil elastase (36, 37). Each of these proteins co-localized to extracellular DNA structures extending from the PMs (Figure 1B). The staining for MET-associated proteins was specific as no fluorescent signal was seen when either a secondary conjugated antibody alone or an isotype control secondary conjugated antibody was used to evaluate these structures (Sup. Figure 1). Together, these data suggest that these structures are METs released by PMs. The extent of MET release was then quantified, and PMs co-cultured with GBS released significantly more METs than uninfected cells and DNase I treatment degraded these extracellular structures (Figure 1C). Additionally, MET release by PMs occurred in a dose dependent fashion (Sup. Fig 2A), and MET release was not GBS strain or bacterial species specific as PMs infected with GBS strain GB037, a capsular type V strain, *Escherichia coli*, or heat killed bacteria resulted in similar MET release (Sup. Figure 3).

**Figure 1:**
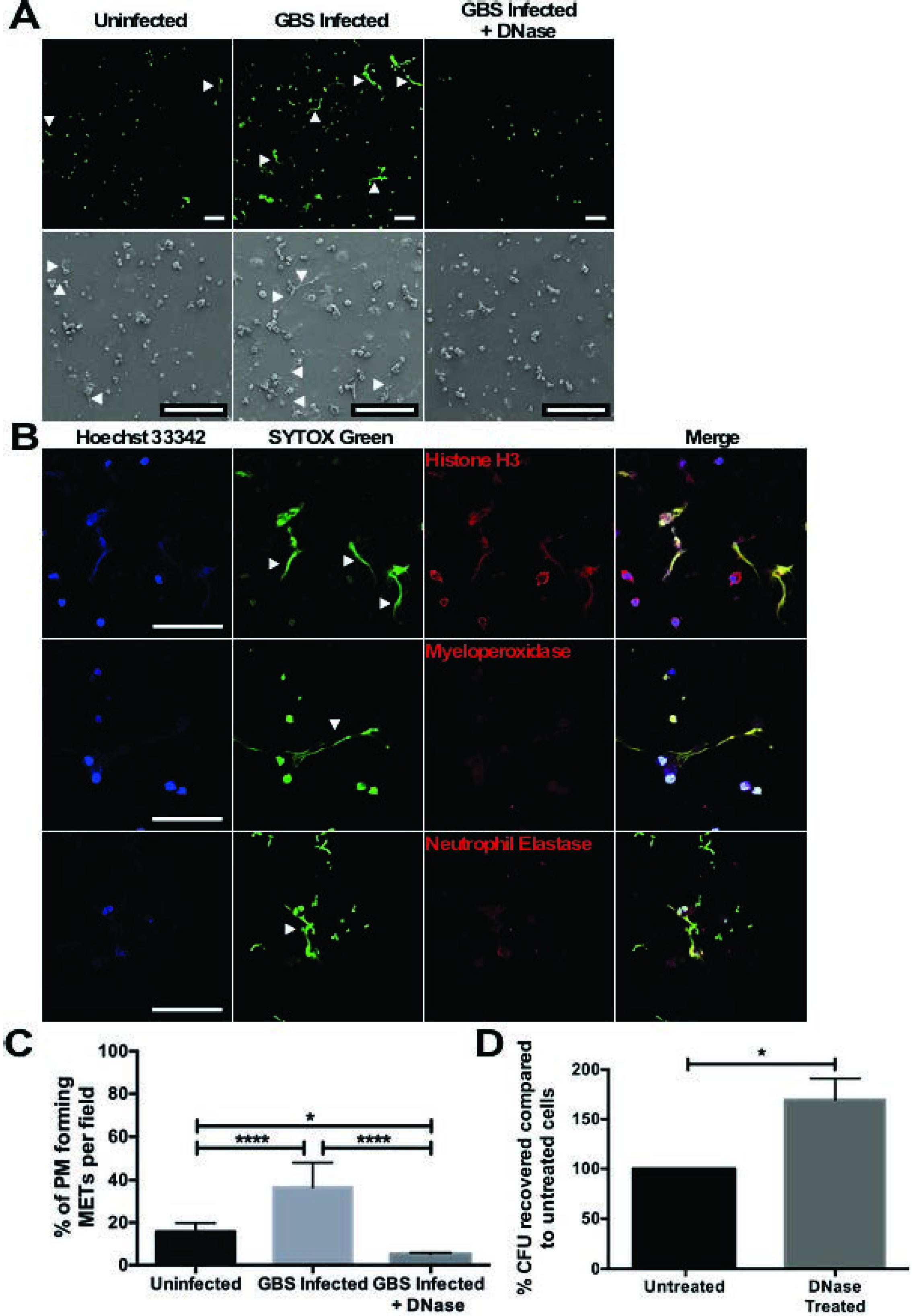
Placental macrophages infected *ex vivo* with GBS release extracellular traps capable of killing GBS cells. 1A: Placental macrophages were infected for 1 hour with GBS cells at an MOI of 20:1. Scanning electron micrographs (bottom row) demonstrate extracellular structures released from macrophages (white arrows), which are not seen after DNase I treatment. PMs were also stained with Sytox Green, a double stranded DNA dye, and evaluated by scanning laser confocal microscopy, which demonstrates extracellular structures composed of DNA (white arrows). Measurement bars represent 100 μm. 1B: Placental macrophage extracellular traps were stained with Hoechst 33342 (blue), a condensed chromatin/nuclear stain, SYTOX Green (green), and specific antibodies for either histone H3, myeloperoxidase or neutrophil elastase as listed (red). Histone, myeloperoxidase and neutrophil elastase staining co-localizes to extracellular DNA staining suggesting that MET structures contain these proteins. Measurement bars represent 100 μm. 1C: PMs releasing METs were quantified by counting MET producing cells seen in SEM images and expressed as the number of macrophages releasing METs per field. GBS infected PMs release significantly more METs than uninfected cells, and DNase I treatment degraded these structures. Data represent samples from 6-8 different placental samples, one-way ANOVA, *F* = 32.7, *p* < 0.0001, with post hoc Tukey’s multiple comparison test. 1D: Placental macrophage METs kill GBS cells. PMs were infected for 1 hour at MOI 20:1 in the presence of DNase I to degrade METs or without (Untreated). Untreated wells were treated with DNase I for the last 10 minutes of infection to break up DNA complexes prior to serial dilution and plating. DNase I treatment significantly impairs bactericidal activity. Data represent the percent recovered colony forming units (CFU), normalized to untreated cells from 7 separate experiments from different placenta samples, Student’s *t* test, *t* = 3.224, df = 6, *p* = 0.0180. **** represents *p* ≤ 0.0001, * represents *p* ≤ 0.05.

One major immunologic function of extracellular traps is the ability to immobilize and kill microorganisms through the locally high concentration of cellular proteins including histones that have antimicrobial effects (37, 38). In order to investigate the bactericidal activity of PM METs, PMs were co-cultured with GBS cells alone or in the presence of DNase I. After 1 hour of infection, significantly more bacterial colony forming units (CFU) were recovered from co-cultures treated with DNase I, suggesting that PM METs have bactericidal activity and eliminating METs with DNase treatment impaired bacterial killing (Figure 1D). To verify that DNase treatment itself did not result in significant PM cell death, thus decreasing bactericidal ability, PMs were incubated with DNase I for one hour prior to washing and stimulating PMs with heat-killed GBS for 24 hours. PM TNF-α release was used as a marker of macrophage viability and function; there was no difference in TNF-α from supernatants of cells treated with DNase compared to untreated cells (Sup. Figure 2B). Additionally, live-dead bacterial staining of PMs infected with GBS demonstrated dead GBS cells adjacent to MET fibers (Sup. Figure 2C). Together, these data provide evidence that PMs release METs in response to bacteria and that these structures are capable of killing GBS cells.

Extracellular trap formation, or etosis, occurs by a cell death pathway distinct from pryoptosis and apoptosis (39). To investigate if GBS infection results in different cell death pathways, GBS infected PMs were assayed for LDH release as a marker of cellular death, TUNEL staining as a marker of apoptosis, and IL-1β release to indicate pryoptosis. At one hour of GBS infection, supernatants of PMs co-cultured with GBS demonstrated an increase in macrophage death, determined by LDH release (Sup. Figure 4A). However, GBS infected PMs did not exhibit a significant difference in IL-1β release or TUNEL positive cells compared to uninfected cells treated with vehicle controls at 1 hour (Sup. Figure 4B-D).

### PMA-differentiated THP-1 macrophage-like cells release METs after direct bacterial contact

Experiments were conducted to determine if MET responses against GBS were specific to PMs or might represent a broader macrophage response. The immortalized monocyte-like cell line, THP-1 cells, was evaluated after differentiation into macrophage-like cells with phorbol 12-myristate 13-acetate (PMA) for 24 hours. THP-1 macrophage-like cells infected with GBS released significantly more METs than uninfected cells, and DNase I treatment degraded the MET structures (Sup. Figure 5). The THP-1 MET response required contact with bacterial cells, as treatment of the macrophage-like cells with sterile filtered bacterial culture supernatant did not stimulate MET release compared to uninfected cells.

### Actin polymerization is required for GBS-induced MET release

Actin polymerization has been shown to be important for MET release (36). A similar role of cytoskeletal changes on GBS-induced MET release in THP-1 macrophage-like cells was examined. Treatment prior to infection with the actin polymerization inhibitor, cytochalasin D, but not nocodazole, which inhibits microtubule polymerization, inhibited MET release compared to GBS infected, untreated cells (Sup. Figure 5). As noted below (and shown in Figure 2B), cytochalasin D also inhibited MET release by human PMs infected with GBS.

**Figure 2:**
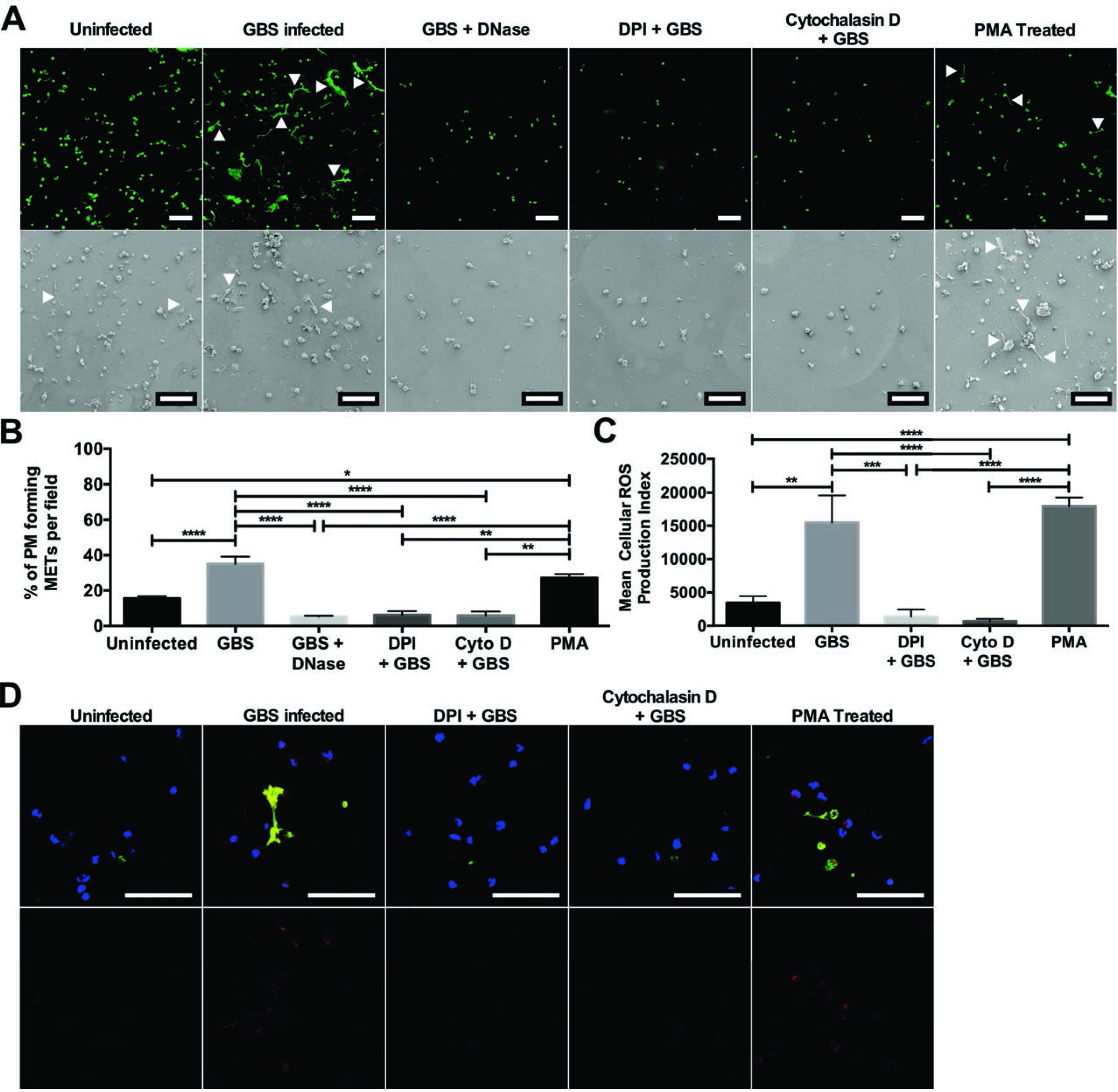
MET release from placental macrophages requires ROS generation. Placental macrophages were incubated with DNase I, DPI to inhibit ROS generation, or cytochalasin D to prevent actin polymerization and then infected with GBS at an MOI of 20:1 for 1 hour. Uninfected cells were also stimulated with 500 nM PMA to stimulate protein kinase C activation. 2A: Cells were then imaged to identify MET release by confocal microscopy after staining with SYTOX green (top row) or via SEM (bottom row). Measurement bars represent 100 μm. 2B: MET release was then quantified as in Figure 1. Treatment with DPI and cytochalasin D significantly inhibited MET release, whereas MET release from PMA stimulated uninfected cells was not different from GBS infected cells. Data represent mean ± SE percent of cells releasing METs per field of 3-9 separate experiments, one-way ANOVA, *F* = 21.1, *p* < 0.0001 with Tukey’s multiple comparison test. 2C, D: PM cell infections were repeated with staining for intracellular ROS production using CellROX deep red reagent. This reagent becomes fluorescent when oxidized by ROS. Cells were co-cultured with GBS cells as above and stained with CellROX deep red, SYTOX green, and Hoechst (2D top row). Measurement bars represent 100 μm. ROS production was quantified by measuring the total red fluorescence per image (2D, bottom row) and the cellular ROS production index was calculated (2C). Data are shown from a representative experiment of 3 independent experiments and are expressed as the mean cellular ROS production index ± SE of 10 images from a single placental sample. GBS infected and PMA stimulated uninfected cells generated significantly higher amounts of ROS than uninfected cells or those treated with DPI or cytochalasin D (one-way ANOVA, *F* = 16.5, *p* < 0.0001 with post hoc Tukey’s multiple comparison test). **** represents *p* ≤ 0.0001, *** represents *p* ≤0.001, ** represents *p* ≤ 0.01, * represents *p* ≤ 0.05.

### Placental macrophage MET responses require ROS production

Neutrophil release of NETs occurs in a ROS-dependent manner (40). It was hypothesized that MET release from PMs may require production of ROS. Treatment of PMs prior to infection with the NADPH oxidase inhibitor diphenyleneiodonium (DPI) inhibited release of METs, whereas treatment of uninfected macrophages with PMA resulted in similar levels of MET release to GBS infected cells (Figure 2A, B). As with the THP-1 macrophage-like cells, treatment of PMs with cytochalasin D prior to infection inhibited MET release. To further define that ROS production was associated with MET release a fluorescent ROS dye was used to evaluate PMs for intracellular ROS production. Treatment with DPI inhibited ROS production, and GBS infection as well as PMA treatment of uninfected PMs resulted in significantly more ROS production than uninfected cells (Figure 2C, D). Interestingly, pretreatment with cytochalasin D decreased levels of intracellular ROS production similar to that of
DPI, suggesting that pretreatment with the actin cytoskeletal inhibitor may actually be preventing MET release by impeding ROS production. Additionally, ROS production in these experiments mirrored the degree of MET release under similar conditions (Figure 2B), suggesting that ROS production is necessary for MET release from these macrophages.

### Placental macrophage METs contain MMPs

During pregnancy, PMs support gestational tissue remodeling through release of MMPs. Because macrophage release of MMPs has been implicated in the pathogenesis of fetal membrane rupture (41), we hypothesized that these proteases may also be released in METs. Five MMPs that have been implicated in development and pathologic pregnancies were evaluated. Immunofluorescent staining of METs was significant for the co-localization of MMP-1, -7, -8, -9, and -12 with extracellular DNA structures (Figure 3A). As MMPs are present within METs and GBS infection induced MET release, metalloprotease concentrations within co-culture supernatants were examined to determine if GBS infection would result in an increase in metalloprotease release. MMP-8 and MMP-9 have been investigated as potential biomarkers for intrauterine infection (42–44), and concentrations of both were significantly elevated in supernatants of GBS infected cells compared to uninfected controls (Figure 3B, C). Global MMP activity of co-culture supernatants was then assessed to determine if the MMPs released were active by using a MMP activity assay, which uses fluorescence resonance energy transfer peptides that, when cleaved by MMPs, are fluorescent. Supernatants taken from placental macrophages co-cultured with GBS demonstrated significantly more MMP activity compared to uninfected controls (Figure 3D). Together these data suggest that PMs express several MMPs, and these MMPs are released during bacterial infection within METs and into the extracellular spaces, where they might contribute to breakdown of gestational tissue extracellular matrix.

**Figure 3:**
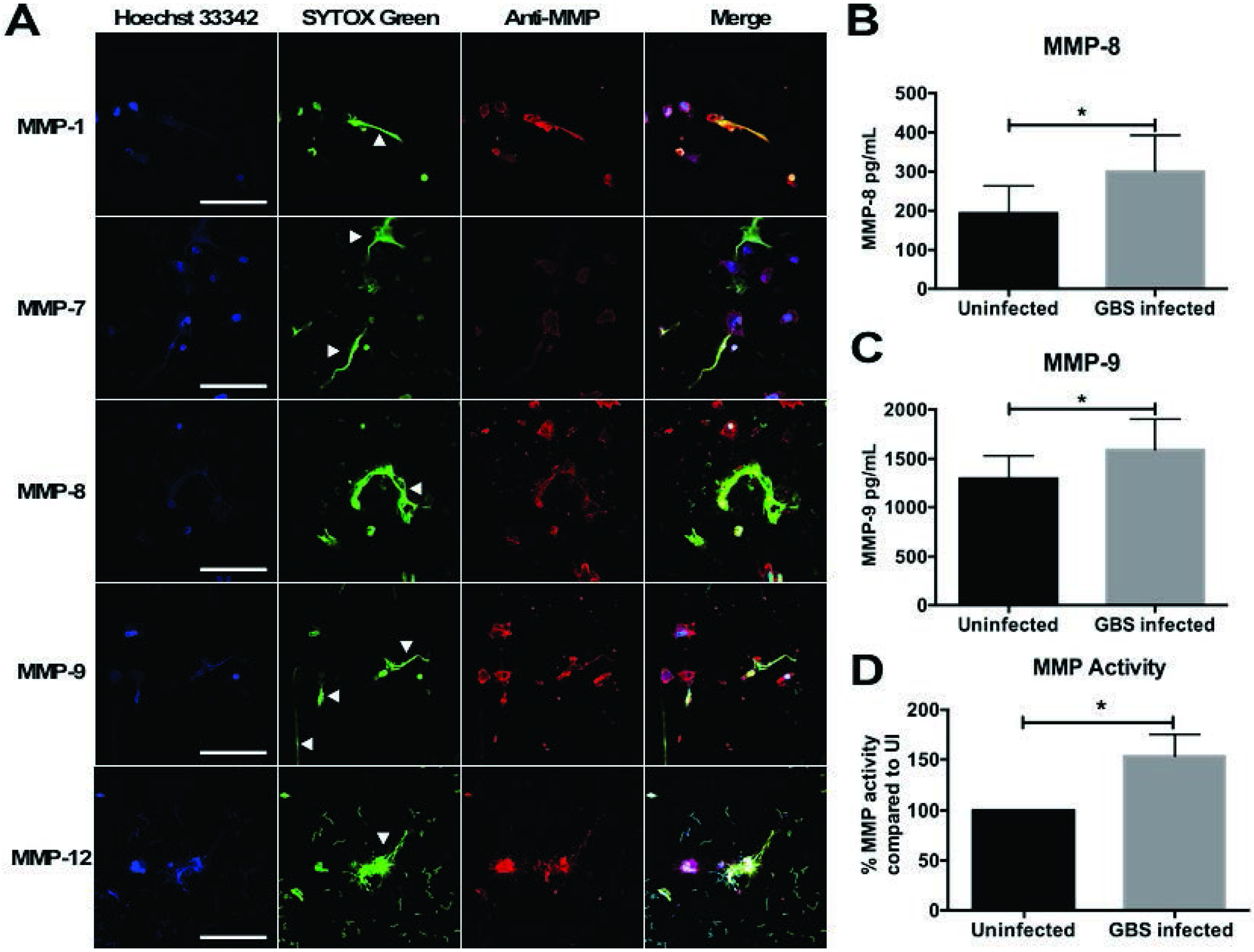
GBS infection results in release of matrix metalloproteinases (MMPs) from placental macrophages. 3A: PM METs contain MMP-1, -7, -8, -9, and -12. PMs were infected as above and then fixed and stained with anti-MMP antibodies and an Alexa Fluor-conjugated secondary antibody (red), Hoechst 33342, and SYTOX green. METs (white arrows) stained strongly for MMPs. Measurement bars represent 100 μm. 3B, C: Supernatant from PM-GBS co-cultures were collected and evaluated for MMP-8 (3B) and MMP-9 (3C) concentrations by ELISA. GBS infection results in a significant increase in MMP-8 (Student’s *t* test, *t* = 3.599, df = 7, *p* = 0.0087) and MMP-9 (Student’s *t* test, *t* = 3.160, df = 10, *p* = 0.0102) release compared to uninfected cells. 3D: PMs release active MMPs in response to GBS. Supernatant from PM-GBS co-culture was evaluated for MMP activity using a MMP Activity Assay to assess global MMP activity within co-culture supernatants. Supernatant from GBS infected cells had 53% more MMP activity compared to uninfected PMs (Student’s *t* test, *t* = 2.439, df = 11, *p* = 0.0329).

### MET structures are present in human fetal membrane tissues infected *ex vivo*

To determine if METs were present within gestational tissues in response to infection, fetal membrane tissues from
healthy, term, non-laboring caesarian sections were obtained, excised, and organized into transwell structures, creating two chambers separated by the fetal membranes. GBS cells were added to the choriodecidual surface and infection was allowed to progress for 48 hours prior to fixing tissues for immunohistochemistry and immunofluorescence analysis. CD163-positive cells were found localized to an area of GBS microcolonies within the membranes that demonstrated histone staining extending beyond the nucleus and into the extracellular space, suggesting the release of a MET-like structure (Figure 4A). Immunofluorescence staining demonstrated CD163-positive cells associated with extracellular material that stained positive for histones and MMP-9 (Figure 4B). Additional staining of fetal membrane tissues using neutrophil elastase as was previously described for identification of NETs in tissues (45), identified cells within the choriodecidua with long extensions that stained strongly for neutrophil elastase that co-localized to histone H3 and DNA staining (Sup. Figure 6). Cells releasing MET-like structures could be compared to cells with intact nuclei and neutrophil elastase staining limited to granule structures suggesting that cells releasing MET-like structures had undergone cellular changes consistent with etosis.

**Figure 4:**
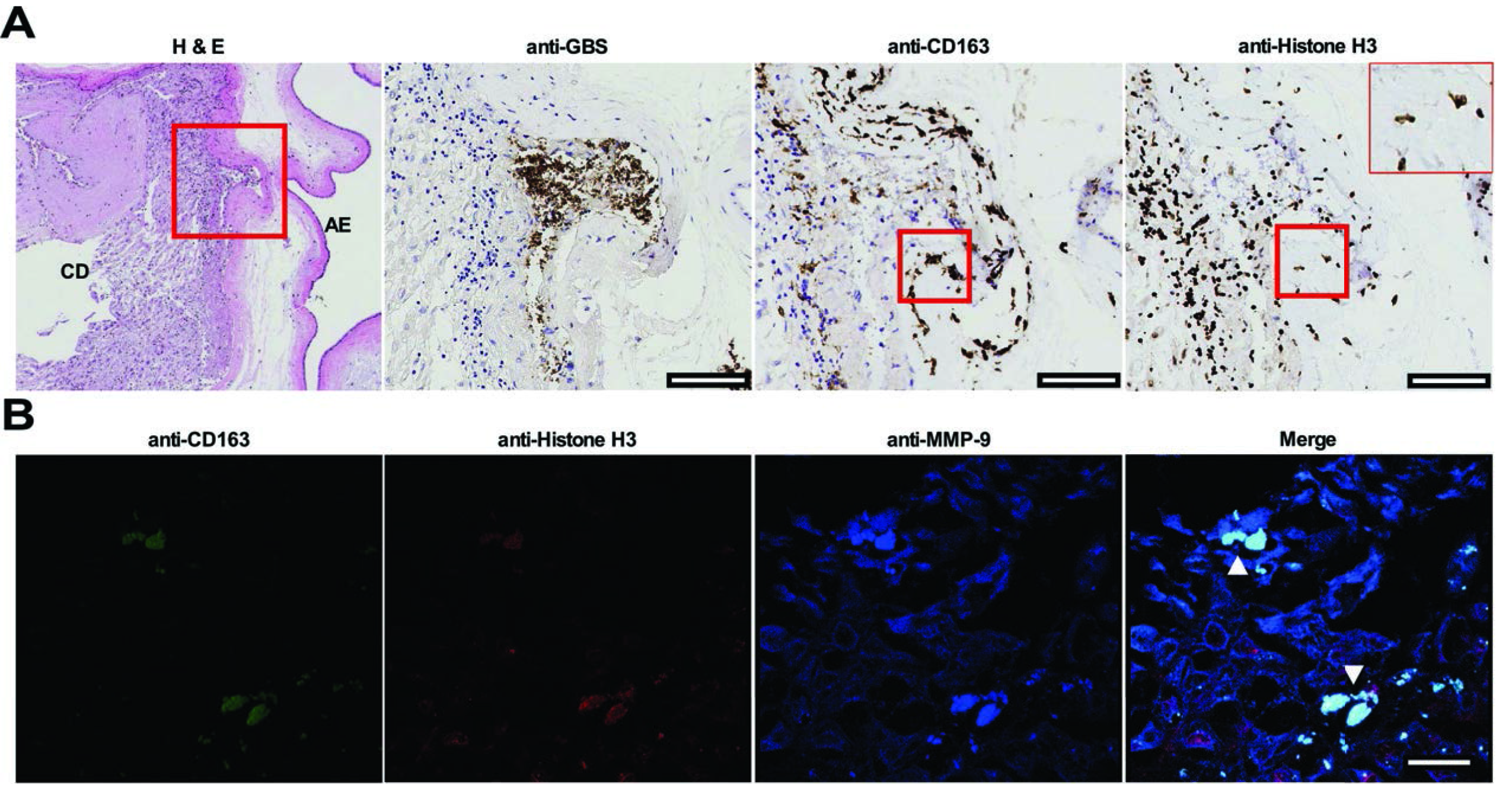
Identification of MET-like structures within human fetal membrane tissues infected with GBS *ex vivo*. Fetal membrane tissues were excised from healthy, term placental tissues from women undergoing routine cesarean section. Fetal membrane tissues were then infected with GBS on the choriodecidual surface for 48 hours prior to fixation and immunohistochemistry (4A) or confocal microscopy (4B). 4A: Fetal membrane tissues were stained with hematoxylin and eosin (far left) and representative images are shown at 4X magnification. Area within the red box is shown in sections stained with anti-GBS, anti-CD163, or anti-histone H3 antibodies and visualized by immunohistochemistry. GBS cells are able to invade from the choriodecidual surface (CD) toward the amnion epithelium (AE). Macrophages are shown in the area of GBS infection and macrophages with extracellular histone staining (far right, 40X insert) are demonstrated in an area that is also staining with the macrophage marker CD163 (red boxes). Measurement bars represent 100 μm. 4B: Fixed and paraffin embedded fetal membranes were stained with conjugated primary antibodies against CD163, histone H3, and MMP-9. CD163-positive cells within the membrane tissue are seen extruding contents that stain positive for histones and MMP-9 consistent with METs (white arrows). Measurement bar represents 20 μm.

## Discussion

In the initial description of NETs in 2004, several potential immune functions were described including the trapping and killing of microorganisms and degradation of bacterial products (37). Release of cellular DNA and proteins within extracellular traps has also been associated with autoimmune pathology in systemic lupus erythematosus and anti-neutrophil cytoplasmic autoantibodies (ANCA)-associated vasculitis, as well as and in diseases of aseptic inflammation (46–50). Other leukocytes including mast cells, eosinophils, basophils and macrophages have now been shown to release extracellular trap structures (36, 51–54).

In this manuscript, PMs are added to a growing list of monocytes and tissue differentiated macrophages capable of releasing METs, which includes human alveolar macrophages, glomerular macrophages, peripheral blood monocytes, and macrophages from other mammalian and non-mammalian species (36). These data demonstrate that human PMs and PMA-differentiated THP-1 macrophage-like cells release METs in response to bacterial infection and after treatment with the protein kinase C agonist, PMA. The present data correlate with previous reports that neutrophils and murine macrophages release traps in response to GBS infection and that METs are capable of killing GBS cells (55, 56).

These data also demonstrate that PM METs contain many proteins previously identified in NETs, including histones, neutrophil elastase, and myeloperoxidase (37). These results mirror other MET investigations demonstrating that diverse macrophages produce and release proteins including neutrophil elastase within MET structures. For example, human glomerular macrophages releasing METs containing myeloperoxidase has been demonstrated in cases of ANCA-associated glomerulonephritis and human alveolar macrophages have been shown to release METs containing histones and MMP-9 (57, 58). Human blood monocytes release METs containing H3 histones, myeloperoxidase, lactoferrin, and neutrophil elastase in response to *Candida albicans* cells, and similar MET contents have been demonstrated in THP-1 macrophage-like cells infected with *Mycobacterium massiliense* (59, 60).

Similar to neutrophil etosis, these data suggest that ROS generation is necessary for PM MET release as this response was inhibited by treatment with the NADPH oxidase inhibitor, DPI. These results mirror reports that inhibition of ROS production via chemical inhibitors resulted in diminished MET release in bovine, caprine, murine, and human macrophages (61–65). In neutrophils, ROS act to break down intracellular membranes and activates neutrophil elastase, which translocates to the nucleus where it degrades histones and promotes chromatin decondensation (39). Myeloperoxidase is thought to contribute chromatin decondensation by a enzymatic-independent mechanism (39). It is unclear at this time if neutrophil elastase and myeloperoxidase perform similar roles in macrophages.

Our data indicate an increase in LDH release during infection, which is consistent with reports that MET release results in cell death (66). Etosis has been noted to be distinct from other cellular death pathways including pryoptosis and apoptosis. At one hour of infection when MET responses were identified, there was no significant difference in TUNEL staining or IL-1β release suggesting that PM METs occur by a distinct pathway. This is notable as previous reports have shown that the GBS toxin β-hemolysin is capable of inducing pryoptosis of macrophages, though in this study infection was allowed to progress for 4 hours, longer than that required for the PM MET response (67). Previous studies have also demonstrated that GBS is capable to inducing macrophage apoptosis, but again this occurred over longer periods of infection than the one hour that was capable of inducing MET responses (68).

Pretreatment of THP-1 macrophage-like cells and PMs with the actin cytoskeletal inhibitor, cytochalasin D, inhibited MET release, but not the microtubule inhibitor, nocodazole. Conflicting reports exist regarding the role of actin polymerization in etosis. Studies evaluating bovine macrophages and THP-1 cells demonstrated a decrease in MET release after cytochalasin treatment, but similar treatment of murine J744A.1 macrophage-like cells, RAW macrophage-like cells, and bovine blood monocytes did not have a significant effect on MET release (60, 61, 64, 69). This collection of conflicting reports mirrors the NET literature. In the original NET description, cytochalasin D prevented cell phagocytosis but not NET release (37). Others have documented NET inhibition with nocodazole or cytochalasin D in response to LPS or enrofloxacin (70, 71). Because of the differential responses, some authors have postulated that phagocytosis may be an important first step towards cell stimulation and ROS generation, and cytoskeletal inhibition may block the initial steps toward MET release. Another possibility is that the pretreatment with cytochalasin D may interrupt trafficking of the NADPH oxidase complex, thus impairing ROS production. NADPH oxidase is a complex of six components, and the cytosolic proteins p40^phox^ and p47^phox^ are known to interact with F-actin; treatment with cytochalasins have been shown to interrupt NADPH complex formation and lead to impaired ROS formation (72, 73). Timing of the cytochalasin treatment is important, as treatment of cells after pre-stimulation with molecules such as LPS, which stimulates NADPH oxidase assembly, may actually increase generation of ROS in these cells (74). In our study, macrophages were pretreated with cytochalasin D and were not stimulated prior to infection. It remains unclear if the conflicting literature with regards to the impact of cytoskeletal inhibition on extracellular traps may be explained by the timing of cytoskeletal inhibition and subsequent effects on ROS production.

PMs were found to produce and release several MMPs within MET structures. During chorioamnionitis, inflammatory mediators lead to the production and release of several metalloproteinases including MMP-1, MMP-7, MMP-8, and MMP-9 (75, 76). MMP-9 is considered to be the major MMP responsible for collagenase activity within the membranes, but many other MMPs are thought to contribute to the processes of membrane weakening (75, 77). This study reinforces and expands previous reports that identified placental leukocytes as being able to secrete MMPs including MMP-1, -7, and -9 (78). Several MMPs have been implicated in preterm birth and pathologic pregnancies. MMP-1 and MMP-9 were found to be elevated in placental tissues of women with preterm births compared to women delivering at full term (79). MMP-1 and neutrophil elastase have been shown to stimulate uterine contractions (80). Interestingly, proteomic comparisons of amniotic fluid from women with premature preterm rupture of membranes demonstrated increases in histones (H3, H4, H2B), myeloperoxidase, neutrophil elastase, and MMP-9 in women with histologic chorioamnionitis and proven intrauterine infection, which likely represents the influx of inflammatory cells into these tissues and potentially release of extracellular traps (81). MMP-12, or macrophage metalloelastase, is a key mediator of the breakdown of elastase and has been shown to be important for spiral artery remodeling during parturition, but to date there are no studies demonstrating changes in MMP-12 release during cases of pathologic pregnancies (82). MMP-12 is better studied in conditions of lung pathology including emphysema, and alveolar macrophages are known to release MMP-12 in METs during infection, suggesting that protease release from leukocytes may contribute to this disease process (65). Analogous to a controlled burn, we speculate that tethering MMPs to MET structures allows the host to control the release of these potent enzymes, thereby limiting their capacity to broadly weaken membrane structure in response to infection.

MET release appears to occur within fetal membrane tissue as demonstrated by our immunohistochemistry and immunofluorescence data. This report adds to the growing relevance of these structures within cases of disease pathology. NETs have previously been identified in placenta tissues from women with pregnancies complicated by systemic lupus erythematosus and preeclampsia (83, 84). NETs were also found in fetal membrane samples of women with spontaneous preterm labor due to acute chorioamnionitis (85). Interestingly, in this report, antibody staining with histone H3 and neutrophil elastase was used to denote NET structures, but given our data, this staining pattern would not have differentiated METs from NETs. Additionally, our group and others have demonstrated that in animal models of vaginal colonization and perinatal infection with GBS, neutrophils traffic to GBS-infected gestational tissues and release NETs containing antimicrobial peptides including lactoferrin as a means to control bacterial growth and invasion (86–88).

In conclusion, we demonstrate that placental macrophages as well as PMA-differentiated THP-1 cells respond to bacterial infection by releasing METs. These MET structures contain proteins similar to NETs, including histones, myeloperoxidase, and neutrophil elastase. MET release from these macrophages can be stimulated in the absence of bacterial cells with PMA and is inhibited by pathways that impair ROS production. Placental macrophage METs contain several MMPs that have been implicated in pathologic pregnancies including premature rupture of membranes. MET structures were identified in human fetal membrane tissue infected *ex vivo*. Together these results suggest that placental macrophages, which are thought to help maintain maternal fetal tolerance and aid in extracellular matrix remodeling, are capable of responding to GBS infection in a way that may trap and kill GBS cells but may also release important mediators of fetal membrane extracellular matrix digestion that could potentially contribute to infection related pathologies including preterm rupture of membrane and preterm birth.

## Materials and Methods

### Placental macrophage isolation and culture

Human placental macrophages (PM) and fetal membrane tissues were isolated from placental tissues from women who delivered healthy infants at full term by cesarean section (without labor). De-identified tissue samples were provided by the Cooperative Human Tissue Network, which is funded by the National Cancer Institute. All tissues were collected in accordance with Vanderbilt University Institutional Review Board (approval 131607). Macrophage isolation occurred as previously described (89); briefly, placental villous tissue was minced followed by digestion with DNase, collagenase, and hyaluronidase (all from Sigma-Aldrich, St. Louis MO). Cells were filtered, centrifuged, and CD14+ cells were isolated using the magnetic MACS Cell Separation system with CD14 microbeads (Miltenyi Biotec, Auburn CA). Cells were incubated in in RPMI 1640 media (ThermoFisher, Waltham MA) with 10% charcoal stripped fetal bovine serum (ThermoFisher) and 1% antibiotic/antimycotic solution (ThermoFisher) overnight at 37°C in 5% carbon dioxide. The following day, PMs were suspended in RPMI media without antibiotic/antimycotic and distributed into polystyrene plates. Cells were incubated for at least 1 hour prior to infection to allow for cell adherence to plate or Poly-L-Lysine coated glass coverslips (Corning, Bedford MA) for microscopy assays.

### THP-1 cell culture

THP-1 cells (ATCC, Manassas VA) were cultured in RPMI 1640 media with 10% charcoal treated FBS and 1% antibiotic/antimycotic media at 37°C in 5% carbon dioxide. 24-48 hours prior to co-culture experiments, cells were treated with 100 nM phorbol 12-myristate 13-acetate (PMA) (Sigma-Aldrich) to induce differentiation to macrophage-like cells. Prior to co-culture experiments, cells were suspended in RPMI media without antibiotic/antimycotic and distributed into polystyrene plates containing Poly-L-Lysine coated glass coverslips and allowed to rest for at least 1 hour prior to infection to promote cell adherence.

### Bacterial Culture

*Streptococcus agalactiae* strain GB590 is a capsular type III, ST-17 strain isolated from a woman with asymptomatic colonization (90), and GB037 is a capsular type V strain obtained from a case of neonatal sepsis (91, 92). *Escherichia coli* serotype 075:H5:K1 is a clinical isolate obtained from a fatal case of neonatal meningitis (93). Bacterial cells were cultured on tryptic soy agar plates supplemented with 5% sheep blood (blood agar plates) at 37°C in ambient air overnight. Bacteria were subcultured from blood agar plates into Todd-Hewitt broth or Luria Broth and incubated under aerobic shaking conditions at 37°C in ambient air to stationary phase. Bacterial supernatant was collected and sterile filtered using a 0.1 μm filter (Millipore Sigma, Burlington MA) and incubated with THP-1 cells at a concentration of 10% volume. Bacterial cells were washed and suspended in phosphate buffered saline (PBS, pH 7.4) and bacterial density was measured spectrophotometrically at an optical density of 600 nm (OD_600_) and bacterial numbers were determined with a coefficient of 1 OD600 = 10^9^ CFU/mL.

### Bacterial-macrophage co-cultures

PMs or PMA-differentiated macrophage-like cells in RPMI without antibiotics were infected with GBS or *E. coli* cells at a multiplicity of infection (MOI) of 20:1 unless otherwise noted. Co-cultured cells were incubated at 37°C in air supplemented to 5% carbon dioxide for 1 hour. As stated, some cells were pretreated with 10 μg/mL cytochalasin D (ThermoFisher), 10 nM nocodazole, 100 Units/mL DNase I, 500 nM PMA, or 10μM diphenyleneiodonium chloride (all from Sigma-Aldrich) for at least 20 minutes prior to infection. At 1 hour, supernatants were collected and cells were fixed with 2.0% paraformaldehyde and 2.5% glutaraldehyde in 0.05 M sodium cacodylate buffer (Electron Microscopy Sciences, Hatfield PA) for at least 12 hours prior to processing for microscopy.

### Field-emission gun scanning electron microscopy

Following treatment and infection as above, macrophages were incubated in 2.0% paraformaldehyde and 2.5% glutaraldehyde in 0.05 M sodium cacodylate buffer for at least 12 hours prior to sequential dehydration with increasing concentrations of ethanol. Samples were dried at the critical point, using a CO_2_ drier (Tousimis, Rockville MD), mounted onto an aluminum stub, and sputter coated with 80/20 gold-palladium. A thin strip of colloidal silver was painted at the sample edge to dissipate sample charging. Samples were imaged with an FEI Quanta 250 field-emission gun scanning electron microscope. Quantification of macrophages producing extracellular traps was determined by evaluating scanning electron micrograph images at 750X magnification and counting total macrophages and those macrophages releasing extracellular traps. Extracellular traps were defined as previously described with typical appearing fibers extending from the cell body into the extracellular space (36).

### Confocal Laser Scanning Microscopy

Co-cultures were completed and macrophages fixed as above. Coverslips were washed once with PBS prior to staining with SYTOX^®^ Green (10 μM final concentration, ThermoFisher) for double stranded DNA (dsDNA), and Hoechst 33342 (5 μM final concentration, ThermoFisher) for condensed chromatin (nuclei). Additional staining for histones and MMPs were accomplished by blocking cells in 1% bovine serum albumin in PBS for 30 minutes at 37°C followed by a 1 hour incubation at 37°C with antibodies for histone H3 (ab5103, abcam, Cambridge, MA), neutrophil elastase (ab68672, abcam), myeloperoxidase (ab9535, abcam), matrix metalloproteinase (MMP)-1 (ab551168, abcam), MMP-7 (ab5706, abcam), MMP-8 (ab81286, abcam), MMP-9 (ab38898, abcam), or MMP-12 (ab137444, abcam). Cells were then washed 3 times with 1% BSA in PBS followed by a 30 minute incubation with an Alexa-594 conjugated goat antirabbit secondary antibody (ThermoFisher) and 2 additional washes with 1% BSA in PBS prior to mounting coverslips onto glass microscope slides with Aqua Poly/Mount (Polysciences Inc, Warrington PA). Macrophages were visualized with a Zeiss LSM 710 META Inverted Laser Scanning Confocal Microscope, and extracellular traps were identified by dsDNA staining that extended into the extracellular environment.

### Bacterial killing by macrophages releasing extracellular traps

PMs were infected with GBS cells at an MOI of 20:1 as described above. As indicated some PMs were incubated with100 U/mL DNase I during infection to degrade extracellular trap structures as has been described previously (37). At the end of 1 hour, DNase I was added to previously untreated wells for 10 minutes to release trapped bacterial cells. Supernatants were collected and PMs were permeabilized with 0.05% Tween-20 in sterile ice-cold water to release intracellular bacteria. Samples were vortexed vigorously, serially diluted, and plated on blood agar plates to enumerate bacterial cells. Untreated PMs were compared to DNase I treated cells and data are expressed as the percent colony forming units (CFU) recovered compared to untreated cells. To further evaluate bacterial killing, PMs were seeded onto coverslips and infected as above. Following infection, cells were stained using the Live/Dead BacLight Bacterial Viability Kit (Invitrogen) prior to confocal laser scanning microscopy.

### LDH cytotoxicity assay

Placental macrophages were incubated in RPMI media without antibiotics or serum and infected as above. Supernatants were collected and centrifuged to pellet cellular debris. Supernatants were analyzed using the Cytotoxicity Detection Kit (Sigma-Aldrich) per manufacturer instructions. Results are expressed as percent toxicity using media without cells as the low control and cells treated with 2% Triton X as a high control. Percent cytotoxicity was calculated using the following equation: cytoxicity (%) = (experimental value-low control)/(high control-low control) × 100.

### Apoptosis Assay

Placental macrophages were incubated in RPMI media without antibiotics and infected as above. Following infection supernatants were removed and cells were fixed with 2.0% paraformaldehyde and 2.5% glutaraldehyde in 0.05 M sodium cacodylate buffer for at least 15 minutes. Click-iT Plus TUNEL Assay with Alexa-Fluor 594 dye (ThermoFisher) was used to identify cells undergoing apoptosis and staining was conducted per manufacturer instructions with additional staining that included Hoechst 33342 to visualize nuclei prior to confocal laser scanning microscopy.

### Macrophage viability assay

To determine if DNase I treatment resulted in alterations in PM viability or function, TNF-α release was used a functional measure. Cells were left untreated or treated with 100 U/mL DNase I for 60 minutes. All cells were then washed and fresh media added prior to stimulation with 150:1 heat-killed GBS cells (incubated at 42°C for 2 hours) for 24 hours. Supernatants were collected and centrifuged to pellet cellular debris before TNF-α release was determined using a DuoSet TNF-α ELISA (R&D Systems) per manufacturer instructions.

### Measurement of intracellular ROS production

Measurements of intracellular ROS production was made by staining cells with CellRox^®^ Deep Red Reagent (ThermoFisher) which measures oxidative stress by producing fluorescence upon oxidation by ROS. PMs were isolated and treated as described above. At the time of infection, a cellular stain mixture containing CellROX Deep Red (5 μM final concentration), SYTOX GREEN, and Hoechst 33342 was added to co-cultures. After 1 hour of infection, cells were washed 3 times with PBS before a 15-minute fixation with 3.7% formaldehyde to preserve CellRox Reagent signal. Coverslips were then mounted onto glass slides and visualized with a visualized with a Zeiss LSM 710 confocal microscope as above. Images obtained were analyzed using Fiji Version 1.0 (94). In order to quantify ROS production, a cellular ROS production index was calculated using the following equation: ((total image intensity – (mean background fluorescence × image area)))/total macrophages counted) × (number of macrophages with ROS production/total macrophages counted). Images capturing only ROS staining (without other stains/channels) were measured to determine the total corrected fluorescence for the total image area. Mean background fluorescence was determined by at least 3 different measurements in areas of the image lacking cellular contents (95). Data are presented as the mean +/− SE ROS cellular production index of 10 images per sample.

### Metalloproteinase ELISA

Supernatants from macrophage-GBS co-cultures were collected and centrifuged as above to remove cellular debris. Supernatants were then evaluated for the concentration of human MMP-8 and MMP-9 using DuoSet ELISA kits (R&D Systems, Minneapolis, MN) per the manufacturer’s protocol and protein levels were calculated from a standard curve.

### Matrix metalloproteinase activity

MMP activity of co-culture supernatants was measured using theMMP Activity Assay Kit (abcam). Supernatants were incubated with assay buffer for 30 minutes and fluorescence signal was measured with a fluorescence microplate reader at an Ex/Em = 490/525 nm. Sample values were normalized to uninfected cells from the same placental sample to calculate a percent change for each placental sample assayed.

### Human fetal membrane infections

Fetal membrane tissue was obtained and cultured as previously described (96). Briefly, fetal membranes were excised from placental tissues. Fetal membrane tissue sections were suspended over a 12 mm Transwell Permeable Support without membrane (Corning) and immobilized using a ¼ inch intraoral elastic band (Ormco, Orange CA) so that the choriodecidua was oriented facing up. Both transwell chambers were incubated with Dulbecco’s modified Eagle’s medium (DMEM), high-glucose, HEPES, no-phenol-red cell culture medium (Gibco, Carlsbad, California) supplemented with 1% fetal bovine serum and PEN-STREP antibiotic/antimycotic mixture (Gibco). Transwells were incubated overnight at 37°C in ambient air containing 5% CO_2_ before media was replaced with DMEM, high-glucose, HEPES, no-phenol-red cell culture medium (lacking the PEN-STREP antibiotic/antimycotic mixture). Bacterial cells were added to the choriodecidual surface of the gestational membranes at a multiplicity of infection of 1 × 10^6^ cells per transwell. Co-cultures were incubated at 37°C in ambient air containing 5% CO_2_ for 48 hours at which time membrane tissues were fixed in 10% neutral buffered formalin prior to paraffin embedding.

### Human fetal membrane immunohistochemistry staining

Tissues were cut to 5 μm sections and multiple sections were placed on each slide for analysis. For immunohistochemistry, slides were deparaffinized and heat induced antigen retrieval was performed on the Bond Max automated IHC stainer (Leica Biosystems, Buffalo Grove IL) using their Epitope Retrieval 2 solution for 5-20 minutes. Slides were incubated with a rabbit polyclonal anti-GBS antibody (abcam, ab78846), rabbit polyclonal anti-histone H3 antibody (abcam, ab8580), or a mouse monoclonal anti-CD163 antibody (MRQ-26, Cell Marque, Rocklin CA) for 1 hour. The Bond Polymer Refine detection system (Leica Biosystems) was used for visualization. Slides were the dehydrated, cleared and coverslipped before light microscopy analysis was performed.

### Human fetal membrane immunofluorescence staining

For immunofluorescence evaluation of METs within fetal membrane tissue, tissues were fixed and sectioned as above. Sections were briefly incubated with xylene to deparrafinize. Tissues were blocked for greater than 1 hour with 10% bovine serum albumins (Sigma-Aldrich) before staining with 1/100 dilutions of mouse monoclonal anti-H3 antibodies conjugated with Alexa Fluor^®^ 647 (ab205729, abcam), rabbit monoclonal anti-CD163 antibodies conjugated with Alexa Fluor^®^ 488 (ab218293, abcam), and mouse monoclonal anti-MMP-9 antibodies conjugated with Alexa Fluor^®^ 405 (NBP-259699AF405, Novus biological, Littleton CO) overnight at room temperature. Additional tissues staining were conducted as previously described (45). Tissues were deparrafinized and then incubated in R universal Epitope Recovery Buffer (Electron Microscopy Sciences, Hatfield PA) at 50°C for 90 minutes. Samples were then rinsed in deionized water three times followed by washing with TRIS-buffered saline (TBS, pH 7.4). Samples were permeablized for 5 minutes with 0.5% Triton X100 in TBS at room temperature followed by 3 washes with TBS. Samples were then blocked with TBS with 10% BSA for 30 minutes prior to incubation with 1:50 dilutions of rabbit poly-colonal anti-neutrophil elastase antibodies (481001, MilliporeSigma, Burlington MA) and mouse monoclonal anti-H3 antibodies conjugated with Alexa Fluor^®^ 647 in blocking buffer at room temperature overnight. The following day, samples were washed in TBS followed by repeat blocking with blocking buffer for 30 minutes at room temperature before incubation with 1/00 dilution of Alexa Fluor^®^ 488 conjugated donkey anti-rabbit IgG (Invitrogen) for 4 hours at room temperature. Samples were then washed and incubated with 5 μM Hoechst 33342 for 30 minutes to stain nuclei. After final washes, slides were dried and coverslipped. Tissues were visualized with a Zeiss LSM 710 META Inverted Laser Scanning Confocal Microscope. Images shown are representative of 4 separate experiments using tissues from different placental samples.

### Statistics

Statistical analysis of MET quantifications was performed using one-way ANOVA with either Tukey’s or Dunnet’s post-hoc correction for multiple comparisons and all reported *p* values are adjusted to account for multiple comparisons. MMP activities assays and bacterial killing assay were normalized to untreated or uninfected cells and analyzed with Student’s *t*-test or one-way ANOVA. *p* values ≤ 0.05 were considered significant. All data analyzed in this work were derived from at least three biological replicates (representing different placental samples). Statistical analyses were performed using GraphPad Prism 6 for MAC OS X Software (Version 6.0g, GraphPad Software Inc., La Jolla CA).

## Acknowledgments

The authors thank Dr. Oscar Gomez-Duarte and Dr. Shannon Manning for providing the clinical bacterial isolates used in this study. The funders of this study had no role in study design, data collection and interpretation, or the decision to submit the work for publication. The authors have no conflicts of interest to disclose. Core Services including use of the Cell Imaging Shared Resource were performed through support from Vanderbilt Institute for Clinical and Translational Research program supported by the National Center for Research Resources, Grant UL1 RR024975-01, and the National Center for Advancing Translational Sciences, Grant 2 UL1 TR000445-06. De-identified, human fetal membrane tissue samples were provided by the Cooperative Human Tissue Network at Vanderbilt University, which is funded by the National Cancer Institute.

---

Figure S1: Staining controls for MET content evaluation. Placental macrophages were treated as in Figure 1B, but were either stained without a primary antibody (top row) or with an isotype control fluorophore-conjugated secondary antibody. Negligible myeloperoxidase (MPO) staining was identified in these samples compared to Figure 1B (middle row) confirming the specificity of the staining protocol. Measurement bars represent 100 μm.

Figure S2: GBS infection of PMs results in release of METs capable of killing GBS. S2A: PMs release METs in a dose dependent response. PMs were infected for one hour at increasing MOI as indicated or treated with vehicle control (PBS) (one-way ANOVA, *F* = 12.3, *p* = 0.0076 with post hoc Tukey’s multiple comparison test). S2B: DNase I treatment does not alter PM viability. PMs were either treated with DNase I or left untreated for one hour before cells were washed and stimulated with heat-killed GBS cells (MOI 150:1) or left unstimulated for 24 hours. Supernatants were assessed for TNF-α release by ELISA as a measure of viability. Treatment of PMs with DNase I did not have a significant effect on TNF-α release (one-way ANOVA, *F* = 7.75, *p* = 0.0016 with post hoc Tukey’s multiple comparison test). S2C: PM METs are capable of killing GBS cells. PM co-cultures were stained with live-dead bacterial staining including Syto9 and propidium iodide. Both dyes stain DNA but propidium iodide (red) is excluded from live cells. Dead GBS cells (red) are shown in close proximity to MET fibers (white arrows). Measurement bar represents 50 μm.

Figure S3: Placental macrophages release extracellular traps in response to different GBS strains as well as *E. coli* cells. S3A: Placental macrophages were co-cultured with live GBS strain GB037, *E. coli* cells, or heat killed GBS or *E. coli* cells at an MOI of 20:1 for 1 hour. Cells were pretreated with DNase I as indicated. Cells were then fixed and subsequently stained with SYTOX Green and evaluated for MET release by confocal microscopy. Measurement bars represent 100 μm. S3B: Placental macrophages releasing METs were quantified by counting MET producing cells from SEM images (not shown) and expressed as the number of macrophages releasing METs per field. At 1 hour of infection live GB037, heat killed GB590 (GBS), and live or dead *E. coli* stimulated MET release as DNase I treatment significantly reduced the number of extracellular structures (unpaired *t*-test of similar treated groups of at least 3 separate experiments from separate placental samples). *** represents *p* ≤ 0.001, ** represents *p* ≤ 0.01, * represents *p* ≤ 0.05.

Figure S4: GBS infection results in PM cell death but not pryoptosis or apoptosis at 1 hour of infection. PMs were isolated and co-cultured with GBS as in Figure 1. S4A: Following 1 hour of infection, co-culture supernatants were assayed for LDH release and percent cytotoxicity was calculated. GBS infection results in a significant increase in cell death (two-tailed, paired Student’s *t* test, *t* = 4.13, df = 4, *p* = 0.0145). S4B: GBS infection does not result in significant PM pryoptosis at 1 hour. Following infection as above, co-culture supernatants were assessed for IL-1β release by ELISA. GBS infection does not result in significant IL-1β release (two-tailed Student’s *t* test, *t* = 0.08945, df = 11, *p* = 0.9303). S4C,D: Following infection as above, PMs underwent TUNEL staining to evaluate cells for apoptotic changes. S4C: Representative confocal images demonstrate nuclear staining (blue) and TUNEL positive cells (red, bottom row). Permeabilized, DNase I treated cells are shown as a positive control. S4D: Quantification of TUNEL positive cells. One hour of GBS infection does not result in an increase in TUNEL positive PMs (two-tailed, paired Student’s *t* test, *t* = 1.056, df = 2, *p* = 0.4017).

Figure S5: PMA activated THP-1 macrophage-like cells release METs in response to GBS. S5A: THP-1 cells were incubated with 100 nM PMA for 24 hours prior to infection to induce differentiation to macrophage-like cells. Cells were infected with GBS at an MOI of 20:1 for 1 hour. As indicated, cells were pre-incubated with DNase I, cytochalasin D, nocodazole, or exposed to 10% volume of sterile filtered bacterial supernatant from GBS cultures grown overnight to steady state. After infection, cells were fixed and evaluated by confocal microscopy after staining with SYTOX Green (top) or by SEM (bottom). White arrows denote METs. Measurement bars represent 100 μm. S5B: Macrophages releasing METs were quantified by counting MET producing cells seen in SEM images and expressed as the number of macrophages releasing METs per field. Data represent mean percent of cells releasing METs per field of 3 separate experiments, one-way ANOVA, F = 8.08, p = 0.028 with Dunnett’s multiple comparison test with samples compared to GBS infected. * represents *p* < 0.05, ** represents *p* < 0.01.

Figure S6: MET-like structures containing neutrophil elastase are seen in human fetal membrane tissues infected *ex vivo* with GBS. Human fetal membrane tissues were isolated and infected as in Figure 4 and then stained for neutrophil elastase (green), histones (red), or DNA/chromatin (blue). Neutrophil elastase positive cells were identified in the choriodecidua (CD) (top panel). The area in the red box was then evaluated at higher magnification and elongated structures of neutrophil elastase that co-localized with staining for histones and DNA consistent with METs were identified (white arrows). This staining pattern contrasts with staining of intact cells where neutrophil elastase staining was isolated to granule structures that did not localize to histone or DNA staining (yellow arrow). Measurement bars represent 20 μm.

